# Building insightful, memory-enriched models to capture long-time biochemical processes from short-time simulations

**DOI:** 10.1101/2022.10.17.512620

**Authors:** Anthony J. Dominic, Thomas Sayer, Siqin Cao, Thomas E. Markland, Xuhui Huang, Andrés Montoya-Castillo

## Abstract

The ability to predict and understand the complex molecular motions occurring over diverse timescales ranging from picoseconds to seconds and even hours occurring in biological systems remains one of the largest challenges to chemical theory. Markov State Models (MSMs), which provide a memoryless description of the transitions between different states of a biochemical system, have provided numerous important physically transparent insights into biological function. However, constructing these models often necessitates performing extremely long molecular simulations to converge the rates. Here we show that by incorporating memory via the time-convolutionless generalized master equation (TCL-GME) one can build a theoretically transparent and physically intuitive memory-enriched model of biochemical processes with up to a three orders of magnitude reduction in the simulation data required while also providing a higher temporal resolution. We derive the conditions under which the TCL-GME provides a more efficient means to capture slow dynamics than MSMs and rigorously prove when the two provide equally valid and efficient descriptions of the slow configurational dynamics. We further introduce a simple averaging procedure that enables our TCL-GME approach to quickly converge and accurately predict long-time dynamics even when parameterized with noisy reference data arising from short trajectories. We illustrate the advantages of the TCL-GME using alanine dipeptide, the human argonaute complex, and FiP35 WW domain.

## I. INTRODUCTION

Biomolecules, such as proteins, dynamically change conformations to perform their functions and thus play a critical role in processes such as protein misfolding and aggregation and protein-ligand recognition. Therefore, investigating biomolecular dynamics is essential for discovering next generation therapeutics, developing novel antibiotic targets, and elucidating protein folding mechanisms that underlie diseases such as Alzheimer’s, Parkinson’s and Cystic Fibrosis.^1^ Indeed, all-atom molecular dynamics (MD) computer simulations can offer insight at resolutions beyond standard experimental setups. However, since small atomic motions such as vibrations occur on the order of femtoseconds, whereas the complex motions at the heart of large conformational changes that drive processes such as protein folding and allostery span timescales from microseconds to seconds, a direct atomistic simulation of such long-timescale motions is only feasible for relatively small biological systems.

Markov state models (MSMs) are a powerful approach that have emerged to tackle this grand challenge.^2–12^ Currently, widely used open-source libraries offer robust implementations for constructing MSMs.^13–15^ MSMs benefit from massive parallelism by exploiting many short molecular dynamics simulations to capture the long-time configurational dynamics that reveal the mechanisms of biomolecular processes.^16^ This is accomplished by partitioning configuration space into a set of states: distinct structures whose component configurations interconvert on a faster timescale than with those belonging to different structures. Identifying the slowest interconverting structures, however, remains a formidable problem.^7,17–25^ This difficulty arises from the fact that, to perform a perfect partitioning, one needs detailed knowledge of the full free energy landscape of a complex condensed phase system. Instead, one is generally limited to a set of states that evolve on slow timescales but are not optimally partitioned.^16,26^ With such a set of configurations, an MSM then provides a discrete-time kinetic description of the interstate conversion, enforcing an effective separation of timescales by requiring transitions between states have no dependence on the history of the system. In this memory-less, or Markovian, limit the rate constants in the kinetic scheme are time-independent. This kinetic description provides an *approximation* to the true dynamics and its accuracy depends on the extent of timescale separation. For a sufficiently accurate (‘valid’) MSM, the maximum resolution in time (minimum time step) allowed by the approximate description is termed the ‘Markovian lag time’. Formally, the intrastate relaxation establishes a *lower bound* to the lag time,^16^ which is the minimum simulation time required for MD data to parameterize the model.

Ultimately, what one would want is a handful of states that provide chemical interpretability for understanding complex biomolecular mechanisms. However, algorithms designed to maximize this timescale separation usually produce many, physically obscure states. This is because downfolding to a biologically intuitive space subsumes slower interstate dynamics of the many-state space into the intrastate dynamics of the reduced space,^27^ increasing the lag time. For example, to model the millisecond folding of the NTL9 peptide using the available simulation data, Pande and coworkers required an MSM containing 2,000 states (with a lag time of 12 ns),^28^ while recent work on the RNA Polymerase (RNAP) II back-tracking necessitated MSMs consisting of 800 states to reach Markovianity within the affordable trajectory.^29^ Therefore there is a balance to be drawn: one wants to coarse-grain aggressively to facilitate interpretability, yet this generally leads to long lag times, which result in both poor temporal resolution and the need to perform longer MD simulations.

Recent work has demonstrated that one can employ non-Markovian theories to resolve the tensions at the heart of the MSM, increasing the resolution to be equal to the MD time step,^30–34^ while simultaneously using only a fraction of the data in the models’ construction.^35^ Of these, the GME, recently used in its time-convolution form as quasi-MSMs (qMSM), provides a particularly useful tool. Indeed, qMSMs have proven useful in tackling important problems such as the gate opening motion of a bacterial RNAP and the mechanism of messenger RNA recognition and inhibition via the RNA-induced silencing complex.^36,37^ Like MSMs, GMEs are most efficient when there is a separation of timescales between intra- and interstate dynamics. Unlike MSMs, GMEs encode the intrastate dynamics into a time-dependent friction function—a memory kernel—removing the approximation of perfect timescale separation. It is this explicit description of the non-Markovianity that allows the improved resolution in time. Yet, their construction lacks conceptual appeal, as they require a convolution integral with the memory kernel which precludes interpretation of the dynamics as a simple kinetic scheme. This motivates the question: is it possible to combine the interpretability of the MSM with the improved accuracy, resolution, and efficiency of GMEs?

In this work, we employ a time-convolutionless (TCL) GME approach that, like the qMSM, encodes the non-Markovian dynamics associated with intrastate motions but, unlike the qMSM, conserves the chemically intuitive nature of MSMs through the action of a generalized non-Markovian rate matrix. We show this easy, accurate, and efficient GME-based approach can capture the biomolecular dynamics of systems of varying complexity, with the resulting dynamics constituting an improvement that combines the advantages, while removing the limitations, of both qMSM and MSM approaches. Indeed, not only does the TCL-GME approach perform just as well as the qMSM on systems that can be exhaustively sampled, but in more difficult cases where all methods struggle to treat statistically underconverged MD data, the TCL-GME can be systematically improved in a manner that has no apparent analogue in the qMSM (or MSM) case. We achieve this through a simple averaging protocol that leverages the onset of Markovian behavior to tame the deleterious effect of noise. Upon reformulating the TCL-GME in discrete-time,^38^ our averaging procedure provides a simple and robust scheme to capture the complex dynamics of biomolecular motions, even in cases that suffer from poor temporal resolution. Finally, in the extreme case where our averaging procedure includes the entire non-Markovian region, our TCL-GME reduces to a high-resolution version of the analogous MSM, recapitulating its identity as the non-Markovian generalization of the conventional MSM and fully elucidating the source of improvement over the traditional time-local approach. We demonstrate that our discrete-time method remains robust even when benchmarked against MD data that extends into the microsecond regime: two orders of magnitude longer than the time required to parameterise the model in question. The strict improvement of our time-local approach is epitomized by its ability to converge an computational sensitive experimental observable (the folding time) using less than half of the data required by the traditional MSM.

## II. CONNECTING MARKOVIAN AND NON-MARKOVIAN EVOLUTION

Whether one wants to directly use a long MD trajectory or many short MD simulations to elucidate complex biomolecular processes, the first task is to find the states that will provide one with the basis of a mechanistic interpretation. The second task is to construct an accurate and efficient description of the dynamics of such configurations. As we mentioned in the Introduction, below we do not consider how one identifies these configurational basins (the interested reader can see, for instance, Refs. 18–21, 23–25, and 39), but rather focus on the second problem: given a set of configurations whose dynamics one can only afford for only short times, how does one construct a dynamical framework to accurately and efficiently capture the dynamics of these configurations over all time?

To characterize the time-dependent transitions connecting states, it is natural to focus on their equilibrium time-correlation functions,

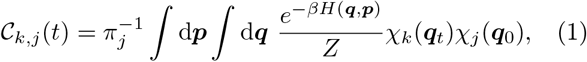

where 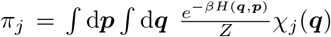 is the equilibrium probability of state *j*, the {*χ_k_*} are mutually orthogonal indicator functions that define the continuous sets of configurations that compose each state, *Z* is the canonical partition function of the system, and ***q*** and ***p*** are the coordinates and momenta of all atoms in the system. Since the states are mutually disjoint, 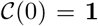. These correlation functions, together the transition probability matrix (TPM), correspond to the conditional probability of finding the biomolecular complex in configuration *k* at time *t* given that it started in configuration *j* at *t* = 0. We now turn to both Markovian (MSMs) and non-Markovian (GMEs) descriptions of the dynamics of the TPM, 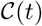.

### A. MSMs and qMSMs

After configuration space has been partitioned into non-overlapping states,^22^ to obtain a valid Markovian description of the TPM dynamics, the MSM framework requires one to identify the smallest time scale *τ_L_* such that the TPM satisfies the Chapman-Kolmogorov condition,^18,40^

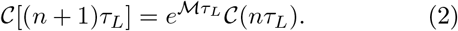

Here, 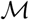 is a time-independent rate matrix and *τ_L_* is defined to be the Markovian lag time. In practice, Eq. 2 is rearranged such that *τ_L_* is found by identifying the onset of a plateau in the implied time scale (ITS), defined as

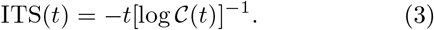

This time scale is associated with the time taken for degrees of freedom within the aggregated states to achieve equilibrium and thus for the systems to become memoryless, or Markovian. Once *τ_L_* is identified, the configurational dynamics can be predicted at integer multiples of *τ_L_*. In other words, *τ_L_* defines the interval at which a given (non-Markovian) biomolecular process can be viewed as Markovian. Consequently, the resulting dynamics are discontinuous,^40^ thus obscuring the observations of dynamical processes which may occur on the interval [*nτ_L_*, (*n* + 1)*τ_L_*]. Furthermore, Eq. 2 implies *τ_L_* sets the lower bound on MD simulation time required to parameterize the MSM that describes 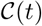.^20^ There is, however, no guarantee that intrastate equilibration will occur within an affordable time scale to perform MD.^16^

Recent work has shown that it is possible to employ a GME approach to account for the effect of memory (non-Markovian) behavior at early times, allowing one to construct a quasi-Markov State Model (qMSM), given by

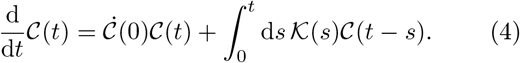

Here, the potentially complex intrastate dynamics are encoded into the time dependent memory kernel 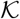.^35^ Crucially, 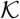 decays to zero on a characteristic time-scale *τ_K_*, termed the kernel cutoff time, enabling one to approximate the upper limit of the integral in Eq. 4 as min {*τ_K_, t*}. It has been further shown that *τ_K_* ≤ *τ_L_*, illustrating that the qMSM approach strictly improves upon the MSM. It does this both by reducing the amount of simulation time needed to capture the exact dynamics, while simultaneously giving access to the dynamical events occurring between multiples of *τ_L_*. Indeed, the qMSM offers remarkable accuracy, temporal resolution, and often requires much less MD simulation time to fully construct the generator of the dynamics, i.e., the memory kernel 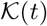.^35^ The qMSM has been profitably applied to, for example, understand the significance of the *β*-lobe of RNA polymerase during transcriptional initiation, ^36^ and elucidate the mechanisms used by the RNA-induced silencing complex to recognize and target mRNA molecules in a sequence specific manner.^41^

Unfortunately, the qMSM is not without its problems. First, evaluation of a convolution integral becomes computationally cumbersome as the dimension of the TPM increases. Second, constructing 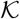 requires the first and second derivatives of 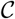,^42^ giving rise to numerical instabilities which we will analyze in a later section. Third, from a qualitative perspective, the qMSM approach obfuscates the physical interpretation of the MSM in terms of “states and rates”. Specifically, the MSM provides a physically intuitive rate matrix, 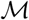, whose diagonals can be interpreted as the likelihood of remaining in a particular state, and whose off-diagonals describe the probability of making a transition from one state to another. In contrast, the memory kernel appears under a convolution integral in the equation of motion for the TPM, Eq. 4, and therefore cannot be understood separately from its cumulative effect over the *history* of the TPM. Hence, the qMSM does not appear to offer a simple way to interpret the memory kernel matrix elements in terms of instantaneous transition rates, e.g., where a number twice as large can be immediately identified as taking half as long to move between two states in a given chemical scheme. These complications motivate the search for an alternative method that accurately and efficiently captures the exact dynamics in a robust, accurate, and intuitive manner.

### B. The TCL-GME

For a non-Markovian theory, such as the qMSM, to be interpreted in terms of rates one would want to write it in a time-local form, comparable to Eq. 2. For this reason, we adopt the time-convolutionless (TCL) GME,^43–45^

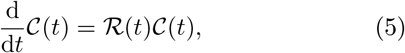

where 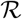 is the time-local generator that encodes the non-Markovian dynamics arising from imperfect timescale separation between intra- and interstate dynamics, and can be understood as a generalized time-dependent rate matrix. Furthermore, the matrix elements of the time-local generator plateau at a characteristic timescale, *τ_R_*,^38,45^ allowing one to separate the time over which non-Markovian evolution takes place (0 ≤ *t* < *τ_R_*) and when Markovian evolution begins,

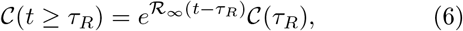

where 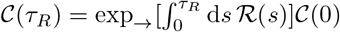 is the value of the TPM at *τ_R_* given by the action of the time-ordered propagator on the initial condition, 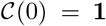, and 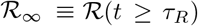 is the long-time limit of the time-local generator. 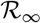 is the time-independent rate matrix that encodes the *true* Markovian evolution of 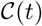 beyond *τ_R_* and elucidates the connection with Eq. 2.

Since the two timescales, *τ_L_* and *τ_R_*, determine the minimal amount of simulation data required to fully construct the MSM and TCL-GME, respectively, it would be profitable to derive a relationship connecting the two quantities. In Appendix A, we analytically demonstrate that

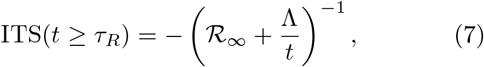

where 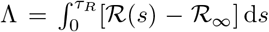 quantifies the deviation that intrastate motions causes on otherwise Markovian interstate transition rates. Comparing this to Eq. 3 allows us to state that

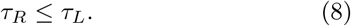

Importantly, Eq. 8 demonstrates that the only cases where an MSM can be as data-efficient as the TCL-GME, albeit at the cost of a lower temporal resolution, is when Λ = 0. This inequality thus enforces a new lower bound on the amount of required simulation time and is one of the central results to the paper, demonstrating that *the TCL-GME always provides a description that is more data-efficient or, at worst, as data-efficient, as the MSM while retaining a high temporal resolution*. What remains to be shown is the relative accuracy and efficiency of the TCL-GME approach in comparison with the qMSM. We will achieve this by comparing the performance of each dynamical approach on three different protein systems of varying levels of complexity: alanine dipeptide,^35^ the human argonaute complex,^41^ and the FiP35 WW domain.^35,46^

## III. ALL-ATOM PROTEIN SYSTEMS

In what follows, we apply the TCL-GME to three systems of varying complexity—alanine dipeptide, argonaute, and FiP35 WW domain—and compare these predicted dynamics to those calculated by both the MSM and qMSM. Here, as previously stated, we do not consider the specifics of how to construct the reduced space but rather restrict our attention to their dynamics. Firstly, for alanine dipeptide we consider a 4 state model with metastable states corresponding to the molecule’s free energy projected onto the backbone torsional angles {*ψ*, *ϕ*}, as constructed in Ref. 35. Secondly, for argonaute, we use another 4 state model from structures corresponding to local minima in the free energy landscape of the first two slowest modes, as constructed in Ref. 41. Finally, for FiP35 WW domain, we use two reduced models: the first contains 3 states and its construction is detailed in Appendix B; the second contains 4 states corresponding to a folded state composed of two *β*-hairpins, an unfolded state, and structures corresponding to both on- and off-pathways, and its construction is outlined in Ref. 35. To clearly benchmark each method while illustrating its advantages and disadvantages, we show only one of the time-dependent conditional probabilities for each protein system. The full time-dependent conditional probability matrices are available in the Supporting Information.

### A. Alanine Dipeptide

We begin our analysis of the TCL-GME and illustrate the utility of the inequality in Eq. 8 using a simple test system, alanine dipeptide. After obtaining TPM dynamics from MD simulation, we construct an MSM as discussed in Ref. 35, and we construct both the qMSM and TCL-GME as described in the Materials and Methods section. In Fig. 1(a) we identify the values of *τ_R_*, *τ_K_*, and *τ_L_* using a root mean square error (RMSE) analysis (see Appendix F) that quantifies the deviation of the dynamics predicted as a function of *τ_L_*, *τ_R_*, and *τ_K_* from the reference dynamics. We use a convergence threshold of 4% of the final value in the RMSE, which leads to graphical accuracy in the resulting dynamics. For the qMSM and the TCL-GME, this leads to *τ_R_* = *τ_K_* = 1.5 ps, while for the MSM the lag time at the same error is *τ_L_* = 10 ps. The results in Fig. 1(b) show the dynamics that would result if one could only use TPM data, obtained from the MD, for the first 1.5 ps; such a choice of *τ_L_* leads the MSM to severely overestimate the equilibration rate. In contrast, Fig. 1(c) shows how a valid MSM is able to capture the exact dynamics, albeit with severely reduced temporal resolution. The drawback of the finite resolution is visible at earlier times, where the (negative) curvature of the MD data is neglected by the MSM but captured by the GMEs. Together, the results of Fig. 1 show that the TCL-GME suffers no loss of performance with respect to the qMSM, with both GMEs able to make accurate high resolution predictions using only 15% of the MD data required to construct a valid MSM.

**FIG. 1.**
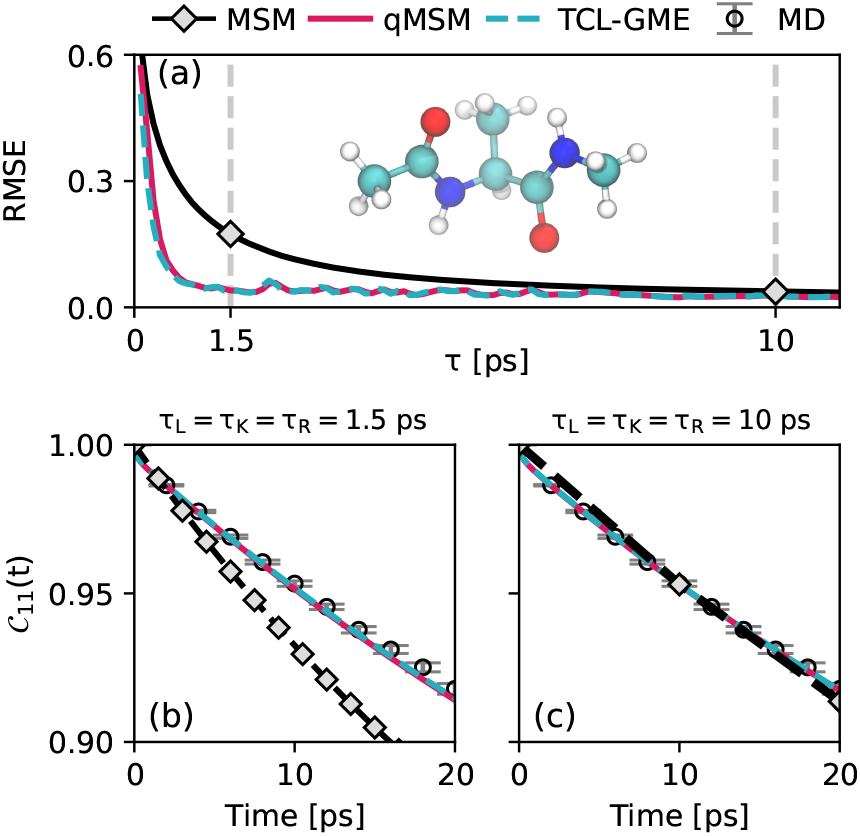
Application of the TCL-GME to alanine dipeptide with comparisons to the MSM and qMSM. (a) Root mean square error (RMSE) curve for the MSM, qMSM, and TCL-GME. Vertical lines show the errors associated with cutoffs (*τ*) of 1.5 ps and 10 ps. Alanine dipeptide is shown (2 residues). (b) TPM dynamics 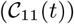 computed with MSM, qMSM, and TCL-GME approaches with *τ_L_* = *τ_K_* = *τ_R_* = 1.5 ps. (c) CK test for 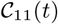 computed with *τ_L_* = *τ_K_* = *τ_R_* = 10 ps. The 4-state TPMs parameterized with *τ_K_* = *τ_R_* = 1.5 ps and *τ_L_* = 10 ps are shown in SI Fig. 1. Error bars were obtained using a bootstrapping approach as shown in Ref. 35.

### B. Argonaute

Will the simplistic form of Eq. 6 maintain a comparable level of performance to the qMSM for a much more complicated system? To address this, we consider the target recognition of human argonaute 2 complex.^37,47^ It is challenging to obtain sufficient MD sampling to model the dynamics of this complex process, which involves coupled conformational changes of messenger RNA, microRNA, and the Argonaut protein. In fact, the ITS curves shown in Fig. 2 do not plateau over the available time window, demonstrating that the available TPM time is not sufficient to construct a valid MSM. That is to say, constructing an MSM is unaffordable at the same level of dimensionality reduction as the faithful qMSM.^41^

**FIG. 2.**
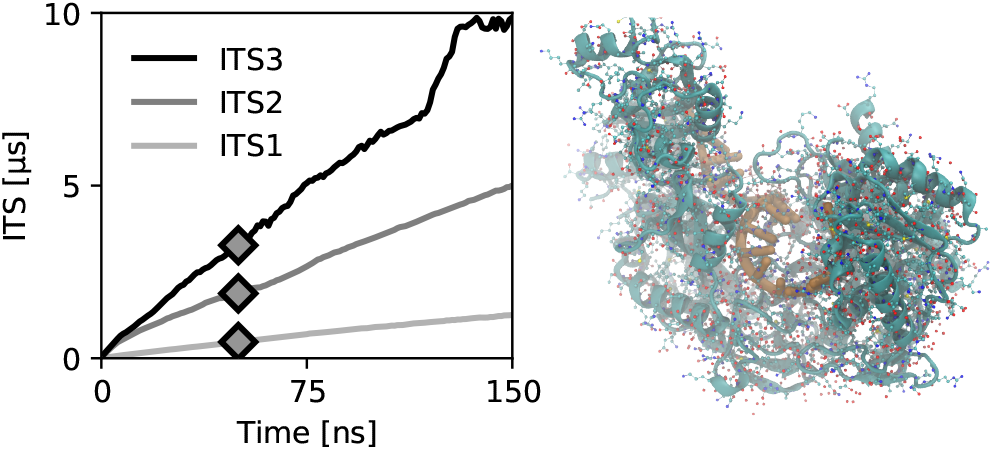
Demonstration that the massive spatial and temporal scales of the argonaute protein present a challenge to MSMs. **Left**: ITS plot of Eq. 3, for the three non-unitary eigenvalues, whose plateau time corresponds to the Markovian lag time, *τ_L_*. Diamonds show the choice of *τ_L_* in Fig. 3, but one can appreciate that no choice for this window of MD data would be satisfactory. Using the 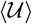-GME approach (discussed in this section), Markovianity is found to require ~ 1200 times as much simulation data. **Right**: Rendering of the argonaute protein containing the mRNA strand used to obtain the MD data. The protein itself is composed of 831 residues.

Owing to the statistical noise that arises from averaging over limited MD data to construct the TPM,^41^ the numerically extracted 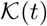 and 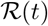 in Fig. 3(a) and (c) also display noise that makes it difficult to graphically identify their cutoff times, *τ_K_* and *τ_R_*, respectively. To illustrate how both GMEs behave as the cutoff time *τ* is increased, we display the dynamics predicted from each method using representative cutoff choices of *τ_R_*, *τ_K_* ∈ {25, 35, 45, 55} ns in Fig. 3(b),(d). Interestingly, the qMSM and TCL-GME perform similarly, with 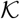 and 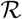 predicting dynamics within the MD error bars using cutoff times of 35 ns. Disappointingly, neither GME exhibits stability with respect to increasing *τ_R_* or *τ_K_*, and the resulting RMSE curves do not monotonically converge towards zero (see SI Fig. 2).

**FIG. 3.**
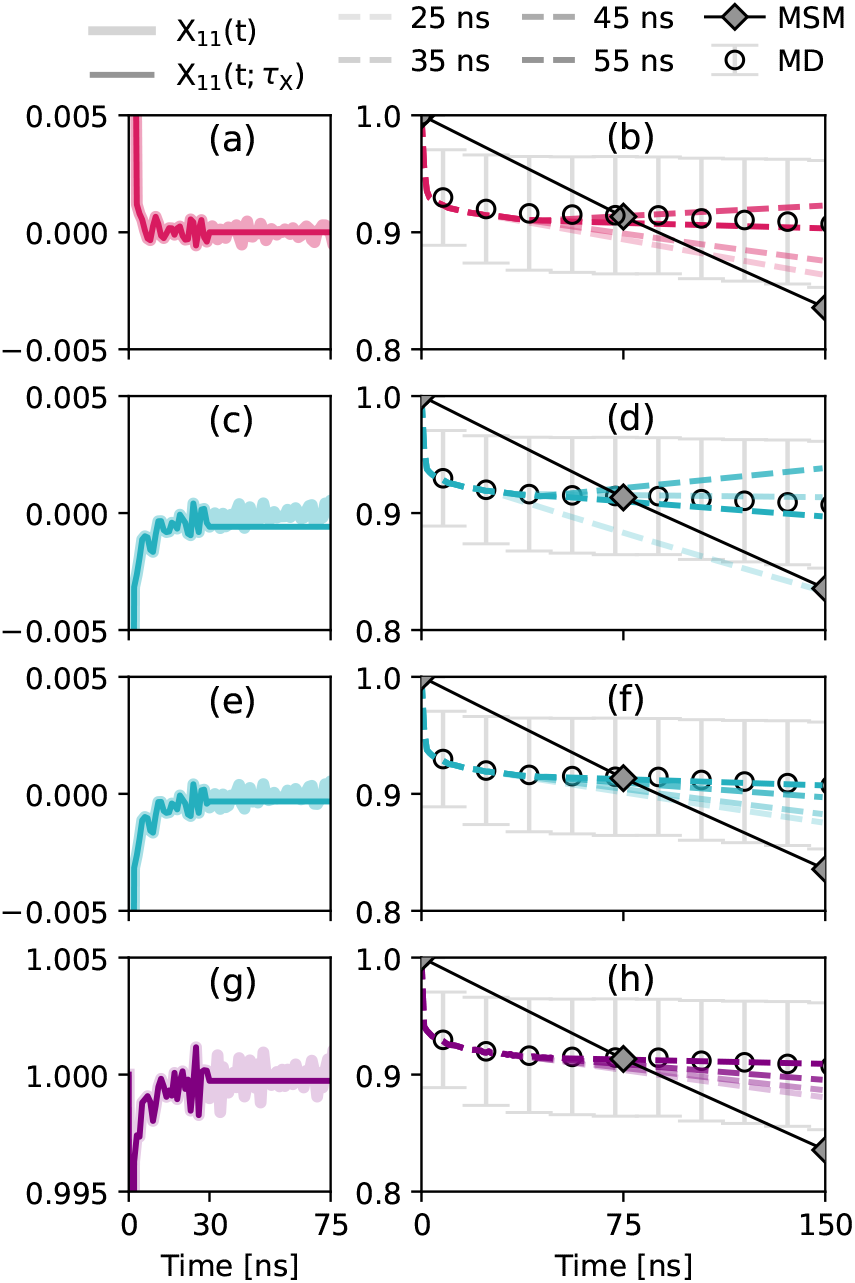
Instability of the qMSM and TC-GME in the case of the Argonaute protein and demonstration of the robustness of our 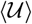-GME approach. (a) The transparent line shows the memory kernel 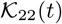 as a function of time. From the RMSE (see SI Fig. 2(a)), we observe that 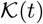 converges by 30 ns. The solid line shows the replacement of 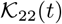 with zero after this time. (b) Dynamics predicted using the qMSM with *τ_K_* ∈ {25, 35, 45, 55} ns, where decreasing transparency corresponds to increasing values of *τ_K_*. (c) Similar to (a), the transparent line shows the time-local generator 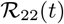 as a function of time, and the solid line shows the replacement of 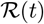 with 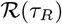 after *τ_R_* = 30 ns. (d) Dynamics predicted using the TCL-GME with *τ_R_* ∈ {25, 35, 45, 55} ns, where decreasing transparency corresponds to increasing values of *τ_R_*. (e) The transparent line shows 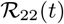 as a function of time, and the solid line shows the replacement of 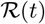 with its average over the window [20, 30] ns after*τ*_R_ = 30 ns. (f) Dynamics predicted using the 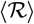-GME. (g) The transparent line shows the propagator 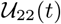 as a function of time, and the solid line shows the replacement of 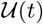 with its average over the window [20, 30] ns after *τ_R_* = 30 ns. (h) Dynamics predicted using the 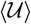-GME. In (b), (d), (f), and (h) we show an MSM parameterized with *τ_L_* = 50 ns. The MD data and error bars were computed using the bootstrapping approach (see Ref. 41 for details).

This lack of controlled convergence can be rationalized by recalling that constructing the GME requires time derivatives of the MD data (See Materials and Methods, Eq. C2). This is true for both 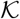 and 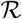. One might hypothesize that the noise in these underconverged MD data is sufficient to compromise the stability of both GME approaches for argonaute. Since TPMs at longer times—like other equilibrium time correlation functions—are constructed from from averaging over less MD data, TPMs at longer times are beset by worse statistical errors.^6,48^ Hence, working with the hypothesis that the fluctuations at later times correspond to noise from statistically under-converged dynamics, we posit a method which averages the noise in 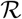 at long times. In fact, during the qMSM approach, truncation at *τ_K_* equates to replacing 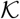 with its long-time average. However, while 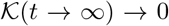 for dissipative problems that equilibrate, we can only estimate it for 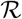.

Visually, Fig. 3(c) suggests that 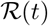 starts to oscillate around its long-time limit around *t* = 10 ns. Thus we introduce an averaging scheme where at *τ_R_* we replace 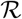 with 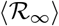, its time average over the interval *[t_r_*, *τ_R_*]. Here, we choose *t_r_* to be the time where the time-local generator appears to have plateaued (See Appendix D). We identify *t_r_* = 10 ns and show the corresponding 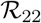 matrix element for *τ_R_* = 30 ns in Fig. 3(e). As Fig. 3(f) shows, with this simple adjustment the TCL-GME converges to the reference dynamics within 55 ns, which strictly improves upon both the MSM and qMSM constructed from the same data. Moreover, the convergence of the TCL-GME with increasing values of *τ_R_* is monotonically decreasing (see SI Fig. 3).

A closer look at Fig. 3(f) reveals that the averaging scheme approaches the reference dynamics from below, but does not actually obtain perfect agreement within these 150 ns. To remedy this, one could average 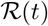 for longer to get a better estimate for 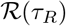. However, this would run counter to our objective of working with the minimal possible MD data. Additionally, as one can appreciate from Eq. 6, error in the estimation of 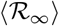 is exponentiated when predicting the GME dynamics. To this end, we propose an alternative route to employ the TCL-GME formalism without requiring any time derivatives or exponentiation of noisy data.^38^ This simply requires re-casting Eq. 6 as

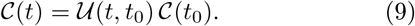

That is, we now work directly with the time-dependent propagator,^49^ 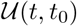, whose construction is detailed in Appendix E. This obviates integration of Eq. 5, and so the noise in the data is never exponentiated during our calculations. Moreover, this method has shown to be robust with respect to low resolution dynamical data in quantum dynamical problems.^38^ Importantly for the protein folding problem, both the time-local interpretability and frugality that result from the plateau at *τ_R_* are unaffected by this manipulation.

Here we extend the protocol proposed in Ref. 38 by combining the direct calculation of 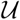 with the aforementioned averaging scheme. This results in our most direct and noise resilient TCL-GME formulation. We identify *t_r_* to be 10 ns and, in Fig. 3(g)–(h), we show the results of this 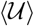-GME. Here, with only minimal adjustments to the original formulation, the 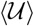-GME monotonically converges to the MD data within 55 ns, maintaining the strict improvements of the TCL-GME over both MSM and qMSM approaches.

With the convergent and stable 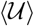-GME dynamics obtained above, we can now determine the true lag time required for a valid MSM description of the dynamics of the 4 states used to elucidate mRNA recognition in the argonaute complex in Ref. 41. To do this, we employ the 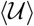-GME to predict the TPM dynamics at long times and use Eq. 3 to obtain obtain the ITS plot (SI Fig. 4). We observe that the ITS curves only plateau by *t* ~ 60 μs, indicating that *τ_L_* is 1200 times larger than the MSM constructed Ref. 41. By comparison the time-local generator cutoff used in our 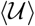-GME, *τ_R_* ~ 50 ns, is more than 3 orders of magnitude smaller, demonstrating that the our approach provides a highly compact and efficient means to fully capture the short-as well as long-time dynamics of complex biomolecular systems.

### C. FiP35 WW-domain

The 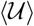-GME method requires two convergence parameters: *t_r_*, the beginning of the averaging window, and *τ_R_*, the total amount of MD simulation time required to parameterize the model (see Appendix E). This begs an important practical question: how does one choose *t_r_* when the onset of the plateau in 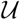 is hidden under the noise? After all, one might expect to observe a lack of convergence when *t_r_* is chosen to be too early. However, by considering a 4-state model of FiP35 WW domain, we find that this is not the case. In this system, where the plateau is not visually obvious (see Fig. 4(c)), we observe that for every choice of *t_r_*, there is a value of *τ_R_* capable of accurately capturing the reference dynamics. In Fig. 4(a) we demonstrate that the *τ_R_* required for the 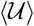-GME to provide accurate dynamics merely increases as *t_r_* is reduced to zero. Indeed, since we know from Eq. 8 that *τ_R_* is bounded above by *τ_L_*, if *t_r_* is given the extreme value of zero then the 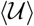-GME reduces to the MSM, with the important distinction that it is able to capture the dynamics between MSM points (see SI Fig. 5; we also give the mathematical justification for this result in Appendix E). In this sense, the 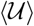-GME parameterized with *t_r_* = 0 constitutes a higher-resolution MSM. The practical implication of this is that while one may make a poor choice of *t_r_* to begin averaging from, one will only pay for this in the length of MD data required to construct the model, *τ_R_*, and not in the final accuracy of the 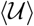-GME dynamics.

**FIG. 4.**
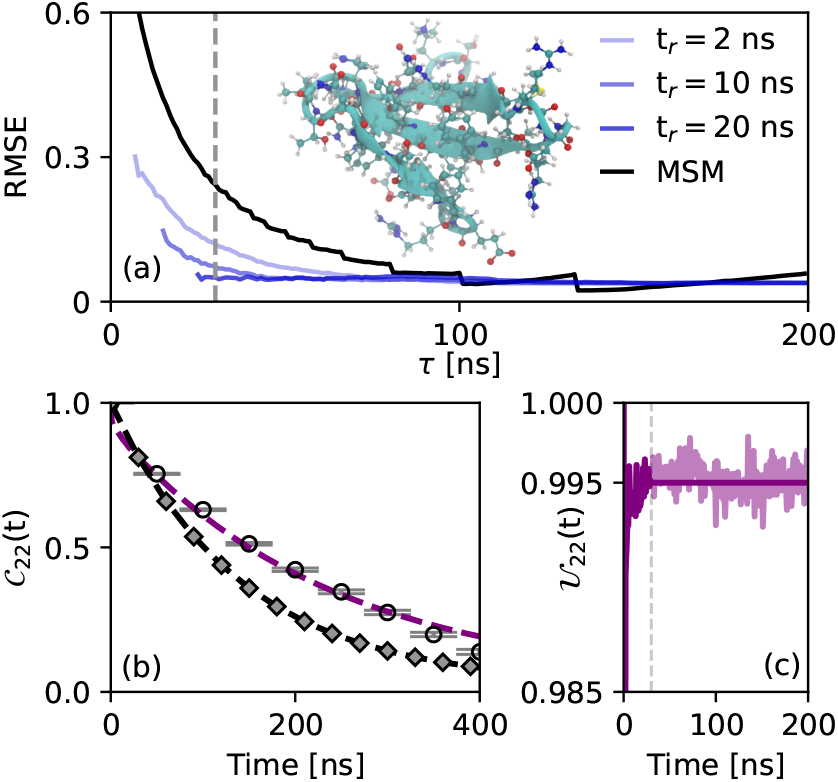
Ability of our 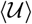-GME to accurately predict the dynamics of the FiP35 WW domain. (a) TPM dynamics 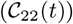 computed using 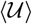-GME and MSM approaches with *τ_R_* = 25 ns (*ℓ* = 5 ns) and *τ_L_* = 25 ns. (b) The propagator 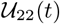 as a function of time, showing that 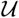 has been replaced with its average at 25 ns. (c) RMSE curves for the MSM and the 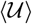-GME as a function of *τ_L_* and *τ_R_*, while varying choices of *t_r_* to illustrate convergence. The structure of the FiP35 WW domain is shown (35 residues).

The best, earliest choice of τR is therefore parametrically dependent on *τ_r_*, but well defined. Since all choices of *t_r_* converge to the same RMSE value, *τ_R_* is robustly identified by a common convergence threshold. To identify the optimal (*t_r_, τ_R_*) pair, we simply find the minimum of the plot of *τ_R_* as a function *t_r_*. Choosing a value of 5% error as converging to the MD dynamics within visual accuracy (see Appendix F), for these FiP35 WW domain data we identify *t_r_* = 20 ns, *τ_R_* = 25 ns, and *τ_L_* = 200 ns, as shown in Fig. 4(a). For comparison, we display the dynamics predicted by both the MSM and 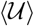-GME when parameterized using only these 30 ns of MD data in Fig. 4(b). In Fig. 4(c), we show the replacement of 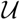 with its average 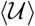 (obtained over the averaging interval of [20, 25] ns). We observe that MSM dynamics predicted using only 25 ns of the MD data set overestimates the equilibration rate, as was the case with alanine dipeptide and the argonaute complex, whereas the 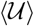-GME parameterized with the same amount of reference data accurately captures the MD data until ~ 375 ns. The small deviation that starts at ~ 375 ns disappears at longer times, where the 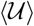-GME correctly captures the long-time limit (see SI Fig. 6). Thus, our analysis shows that accurate predictions of the dynamics from the 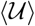-GME require only 15% of the MD data needed to construct a valid MSM.

We now consider the ability of the 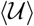-GME to capture the long-time dynamics through a different, experimentally accessible measure: the folding time of the protein. For this, we will consider a 3-state model of FiP35 WW domain (for construction details, see Appendix B) with states one, two, and three corresponding to misfolded, unfolded, and folded structures of the protein, respectively.^46^ Here, we compute the folding time using the mean first passage time (MFPT) procedure outlined in the Materials and Methods section. First, we use the reference dynamics to compute the folding time to be *τ_ref_* = 18.65 μs (SI Fig. 8), which is taken to be the *exact* result for this model, which is in reasonable agreement with the experimentally measured value of 14 ± 1.5 μs.^50^ In particular, if the clustering algorithm does not correctly identify configurations with the folded, unfolded, and misfolded states, this may cause the folding time to appear artificially long. Hence, we focus not on the deviation from the experimental value but rather on the internal consistency between the reference dynamics and the predictions from the 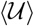-GME and the MSM approaches. To obtain the 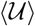-GME predictions of the MFPT, we first identify *t_r_* = 50 μs. As described in Appendix G, we compute the MFPTs corresponding to increasing values of *τ_R_* and *τ_L_* and observe that both the 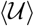-GME and MSM approaches converge to the reference result at long times (see SI Fig.8). We also find that the MSM continuously underestimates *τ_ref_* and appears to continue increasing at times beyond 1000 ns (see SI Fig. 8). In contrast, the 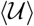-GME remains within 8% of the reference value for the duration of available MD data. Indeed, to converge within 5% error, the 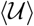-GME requires data up to 168 ns, whereas the MSM does not reach this threshold until 452 ns, suggesting that the 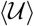-GME provides, even in the estimation of folding times, a more efficient means to capture the long-time dynamics of complex biomolecular systems.

## IV. CONCLUSION

In this work, we have developed and applied a formally exact, chemically intuitive, and systematically improvable approach to modeling non-Markovian biomolecular dynamics. While previous work had exploited the memory of the MSM’s intrastate motions to construct an exact qMSM that could significantly reduce the computational cost required to efficiently predict protein dynamics at long times, it eluded a simple and intuitive chemical interpretation and, as we show here, is highly sensitive to statistical noise in the reference TPM dynamics from which it must be constructed. Here, we have abandoned the time-nonlocal qMSM by moving to a time-convolutionless formulation which admits a simple formal integration, elucidating the analytical connection between GMEs and MSMs and permits a simple interpretation. In particular, not only does this allow the time-local generator to be interpreted as a time-dependent rate matrix, it also allows for systematic improvement in regimes of noisy data. Specifically, we have identified that for cases where the reference TPM suffers from statistical noise (e.g., the argonaute system), a straightforward averaging scheme allows our time-convolutionless approaches (both 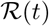 and 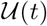) to uniformly converge to the reference dynamics. In contrast, the time-nonlocal approach displays instabilities with increasing simulation time that have no comparable solution without resorting to manipulations of the qMSM formalism.^51^ Furthermore, using alanine dipeptide, FiP35 WW Domain, and argonaute, we have demonstrated that the time-local GME can accurately and efficiently capture short-, intermediate-, and long-time dynamics with no loss of performance. Not only does this approach require an equivalent amount of data as the qMSM, the 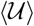-GME requires minimal numerical and physical complexity by eliminating the need for both time-convolution integrals and numerical time derivatives of potentially noisy data. By providing a theoretically robust and physically transparent method to capture the non-Markovian dynamics of a given set of states, we expect the 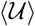-GME to provide a robust scaffold to construct novel methods to find optimal configuration clusters and offer a framework to investigate the mechanisms of complex biomolecular conformational changes.

## ACKNOWLEDGMENTS

A.M.C. acknowledges the start-up funds from the University of Colorado, Boulder. X.H. acknowledges the Hirschfelder Professorship Fund. This work was supported by the U.S. Department of Energy, Office of Science, Office of Basic Energy Sciences (DE-SC0020203 to T.E.M.).

## Appendix A Rigorous Connection of MSM with TCL-GME

Here, we derive Eq. 7 and Eq. 8 from the main text, which rigorously connect the MSM to the TCL-GME. We begin by considering some time t that is strictly greater than *τ_R_* and re-writing 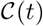 as

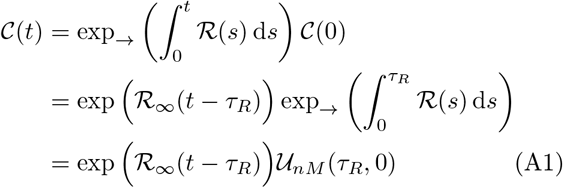

where we have used the fact that the initial condition is the identity matrix, 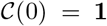 and have introduced *U_nM_*(*τ_R_*, 0) as the propagator over the non-Markovian region. This is equivalent to Eq. 6 in the main text. We insert the above result into the implied time scale equation, defined in Eq. 3, to obtain the result in the main text,

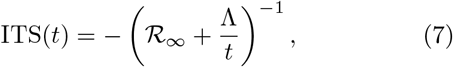

where

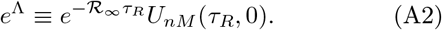

Equation Eq. A2 is exact and easy to calculate given the framework presented here for obtaining the non-Markovian propagator; it can be interpreted as the total deviation in the propagation due to non-Markovian behavior. Keeping in mind that the MSM lag time is taken to be the minimum time-scale associated with the onset of a plateau in an ITS plot, we see see that the right-hand-side of Eq. 7 does not necessarily stabilize for times immediately after *τ_R_*. This allows us to conclude the inequality presented in the main text, that

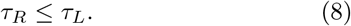

To further simplify its interpretation, one can neglect the effect of time-ordering in the definition of the non-Markovian propagator, which yields the following, modified expression for Λ,

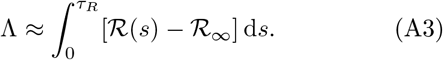

Here, it is clear that Λ approximately corresponds to the integral deviation between the time-local generator over its non-Markovian variation, and its long-time limit.

## Appendix B TPM Construction

The TPM is computed from the transition count matrix (TCM). We first computed the TCM from the MD trajectories. For each lag time *τ*, the raw TCMs (*T^raw^*) were first counted from transition pairs between frames at *t* and 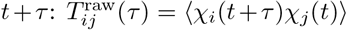, where *χ_i_*(*t*) is the indicator function that determines whether the frames at time *t* is in state *i*. Here, *t* = 0, Δ*t*, 2Δ*t*,…,(*N_traj_* – 1)Δ*t*–*τ*, where Δ*t* is the saving interval of trajectories, and *N_traj_* is the length of trajectories. Normally, detailed balance requires that the TCM be symmetric, i.e., *T_ij_* = *T_ji_*. However, since the raw TCMs are normally not symmetric, we further symmetrize the raw TCMs to satisfy the detailed balance: 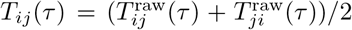.^5^ Finally, we calculated TPMs by column-normalizing the TCM: 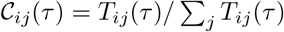.

In our TPM construction, the raw TCM was directly counted from the macrostate models. The four state model of the Alanine dipeptide was constructed with a splitting-and-lumping approach. We first split all the available MD conformations into 1000 microstates using the K-Centers clustering algorithm.^52–54^ Then we lumped the 1000 microstates into 4 macrostates via the PCCA+ (Perron Cluster Cluster Analysis),^55,56^ with the lagtime of 2 ps.

We constructed the three state model of the FiP35 WW domain using tICA (Time-lagged Independent Component Analysis),^57,58^ K-Centers clustering,^52–54^ and PCCA+ (Perron Cluster Cluster Analysis)^55,56^ lumping from MD trajectories provided by D. E. Shaw research. We first performed tICA analysis with pairwise distances between all *α* carbon atoms of the peptide with a lag time of 10ns. Then we used the K-Centers algorithm to generate a 1000-state model based on the top three tICs (Time-lagged Independent Components) of tICA. Finally, we performed the PCCA+ clustering to generate the three state model based on the 1000-state TPM computed at the lagtime of 10 ns.

We constructed the four state model of the Argonaut using spectral oASIS,^59^ tICA, APLoD clustering and PCCA.^55,60^ We employed spectral oASIS to reduce the number of input features, followed by tICA for the dimensionality reduction. Then we grouped the conformations into 81 clusters from the APLoD clustering algorithm, based on the top 4 tICs from the tICA. Finally, we used the PCCA+ algorithm to group the microstates into four macrostates.

## Appendix C qMSM Construction

To solve the integro-differential equation in Eq. 4, we must first construct the memory kernel, 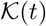, as a function of time directly from the TPM data. We follow Ref. 42 and derive the classical analogue of the self-consistent expansion of the memory kernel

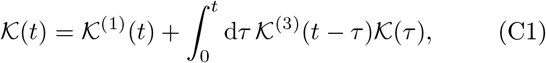

where

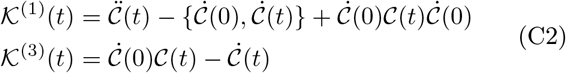

are the projection-free auxiliary kernels.

To compute both 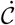 and 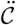, and to thus compute 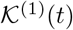 and 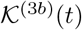, we numerically differentiate the TPM data, 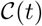. With these auxiliary kernels, we compute 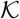 according to Eq. C1 using the discretization procedure in Ref. 61. For completeness, we summarize the algorithm. At the initial time and first timestep, 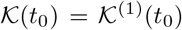 and

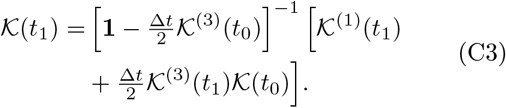

For all subsequent times (*n* ≥ 2),

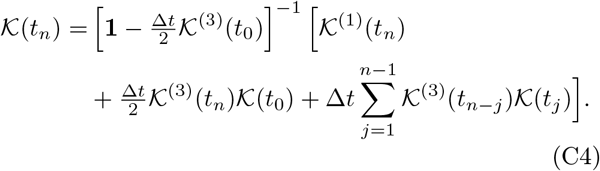

Here **1** is the identity matrix and we employ equally spaced time intervals, such that Δ*t* ≡ *t*_*j*+1_ – *t_j_*.

Once we construct 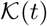, we employ Heun’s method (second-order accurate with respect to Δ_t_) to integrate *Eq*. 4 and obtain 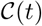. Then we identify an *appropriate* memory kernel cutoff time, *τ_K_*, by applying the RMSE analysis in Appendix C. We approximate the upper limit of the integral in Eq. 4 with *τ_K_*, enabling us to predict the dynamics for times beyond the duration of the MD simulation.

## Appendix D TCL-GME Construction

To reap the benefits of the time-local formalism, we first calculate 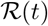 from the TPMs obtained from MD simulation. We do this by rearranging Eq. 5 via matrix inversion to obtain

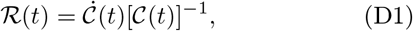

where we calculate 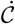 by numerically differentiating the TPM data. As we have discussed, the matrix elements of 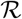 plateau on a timescale, *τ_R_*, associated with the conclusion of non-Markovian evolution, allowing us to set 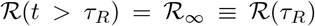. With this definition, we can describe the dynamics after the onset of Markovian evolution, as shown in Eq. 6.

Once we find 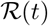, we employ Heun’s method to integrate Eq. 5 and obtain 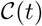. Similar to the discussion in Appendix C, we identify an *appropriate* generator cutoff time, *τ_R_*, using the RMSE analysis discussed in Appendix F.

## Appendix E 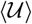-GME Construction

We first formally integrate the TCL-GME in Eq. 5 to obtain

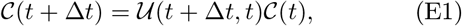

where we have defined 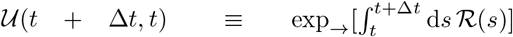 with the “+” subscript denoting the chronological time-ordering of the exponential, as above. We then compute the value of 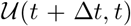 through direct matrix inversion

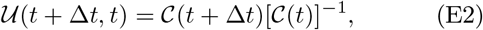

as introduced in Ref. 38. Because 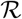 becomes constant at *τ_R_*, the propagator 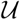 also becomes a constant. Hence, we define 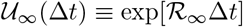. We compute the dynamics beyond *τ_R_* according to

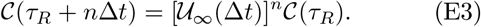

As discussed in our analysis of the Argonaute complex, we developed and implemented a simple averaging scheme capable of taming noise arising from statistically underconverged MD estimates of the TPM. We begin by applying the RMSE stability analysis in Appendix F to determine a valid generator cutoff time; here, we denote this cutoff by *t_r_*. We then introduce another parameter *ℓ* that represents the number of *high quality* TPMs after *t_r_* and denote the corresponding time as *t*_*r*+*ℓ*_. This number is, of course, limited by data availability. To predict the dynamics beyond *t*_*r*+*ℓ*_, we compute the time average of 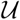 on the time interval [*t_r_*, *t*_*r*+*ℓ*_] using

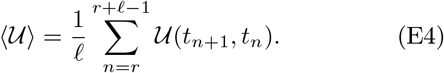

Because our 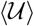-GME requires at least *r* + *ℓ* data points to circumvent the instabilities imposed by noise in biomolecular systems, we generalize our the definition of the generator cutoff time to be *τ_R_* = *t*_*r*+*ℓ*_, *representing not the generator cutoff but rather the minimum amount of data needed to accurately predict the true TPM dynamics*. Ultimately, we recommend that the user performs a rigorous stability analysis with respect to the choices of *r* and *ℓ*.

It can be seen by equating expressions Eq. 6 and Eq. 2 given the same first time step (*τ* = *τ_L_* = *τ_R_*),

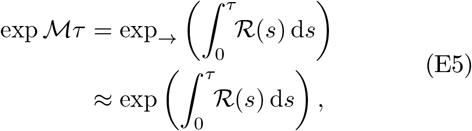

where the right-hand side of Eq. E5 uses the explicit form of the propagator (see Appendix A for details). If this time-ordering of the exponential can be neglected, then we can identify 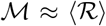. The practical implication of this is that, if we can replace 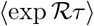 with 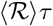 in the 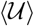-GME, we will obtain exact agreement with the MSM parameterized by the same *τ* (at integer multiples of *τ*). The requirement for this to be true is that 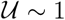, which we show to be satisfied by panel (c) of Fig. 4. Since 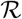 and therefore 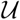 are formally exact before cutoff (by construction they return the reference dynamics),^38^ the dynamics between these MSM points is also accessible to the 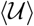-GME. This explains why Fig. 4(c) shows that lim_*t*_*r*_ → 0_(*τ_R_*) = *τ_L_*.

## Appendix F RMSE Analysis

To determine values of *τ_x_* ∈ {*τ_L_*, *τ_R_*, *τ_K_*}, we find the lowest time by which the time-averaged root mean squared error (RMSE), given by

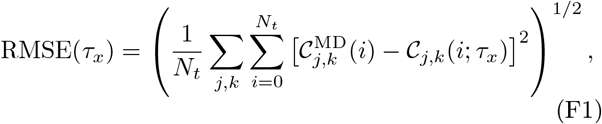

becomes and stays sufficiently small. We identify *τ_x_* to be this minimum amount of time. How small the RMSE should be for any particular application is a choice for the user to determine. In our results, we choose the RMSE to be ~ 5%, which results in graphical agreement between the reference and GME or MSM dynamics. In Eq. (F1), *N_t_* is the number of time steps in the data set.

## Appendix G MFPT method

We apply our newly developed 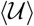-GME to compute folding times for FiP35 WW Domain. To do so, we consider a three state model where states one, two, and three are defined to be the misfolded, folded, and unfolded structures, respectively. To employ Meyer’s mean-first passage time (MFPT) method,^62,63^ we construct the time-dependent MFPT matrix, *M*, as

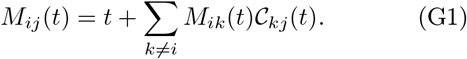

The element *M*_32_ then corresponds to the folding time in this problem.

Practically, one solves Eq. G1 as a system of linear equations.^64^ To solve for the MFPT corresponding to passage to state 3, the folded state, we consider the row 3 MFPT matrix elements and obtain the following system of equations

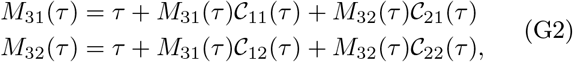

We recast the system in terms of matrices and obtain the final form by matrix inversion,

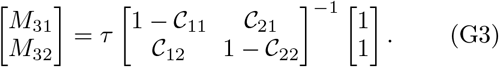

As the dynamics approach equilibrium, the inverse matrix on the right-hand-side of Eq. G3 becomes constant. In practice, we define the folding time to be when *M*_32_/*τ* is within 5% of *M*_32_(*τ*_final_)/*τ*_final_ for the rest of time.

## SUPPORTING INFORMATION

**FIG. 5.**
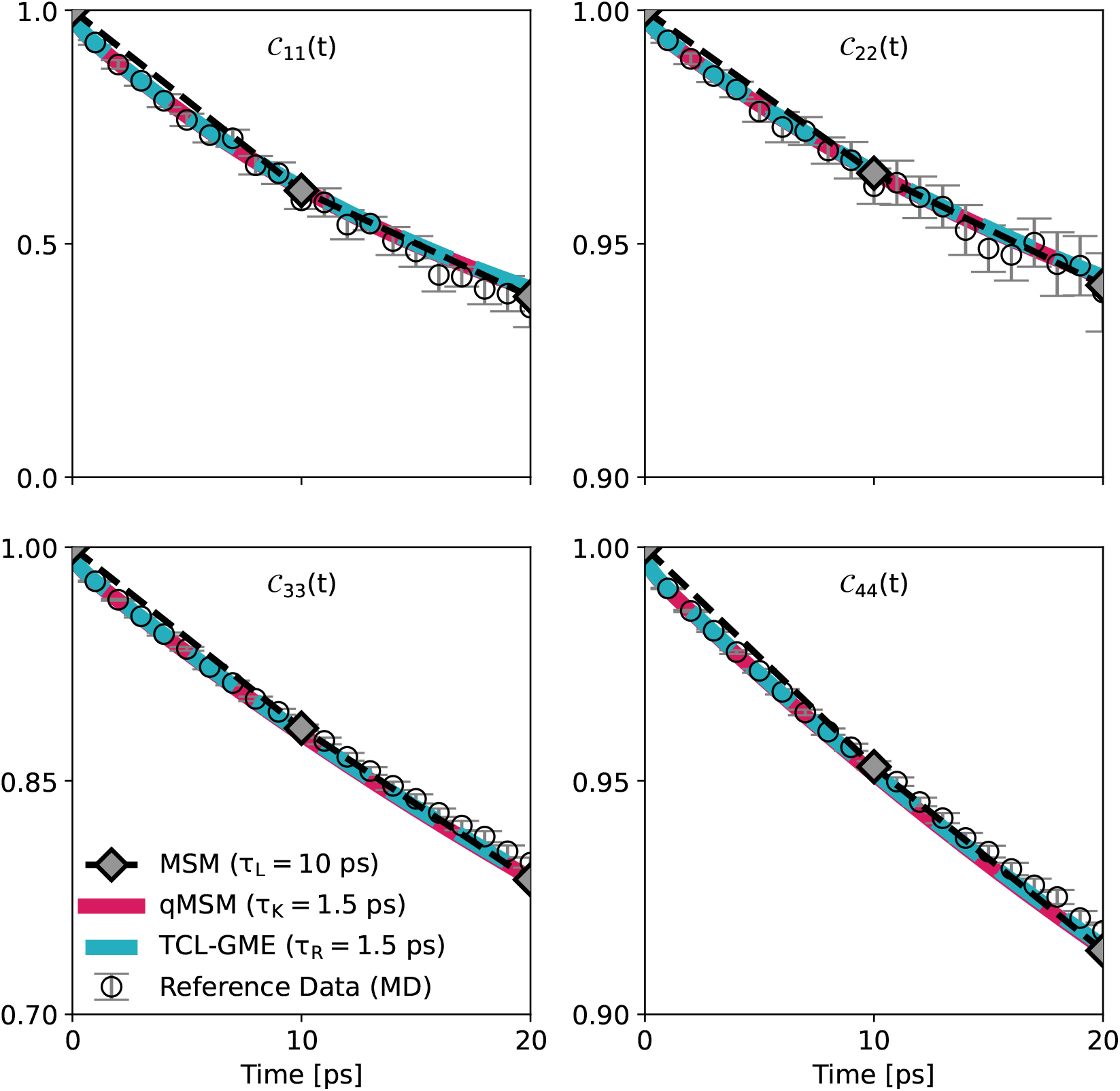
Predicting the 4-state TPM dynamics of alanine dipeptide with MSM (*τ_L_* = 10 ps), qMSM (*τ_K_* = 1.5 ps), and TCL-GME (*τ_R_* = 1.5 ps) approaches, illustrating that the GMEs require 1.5%of the reference data needed to construct an accurate MSM.

**FIG. 6.**
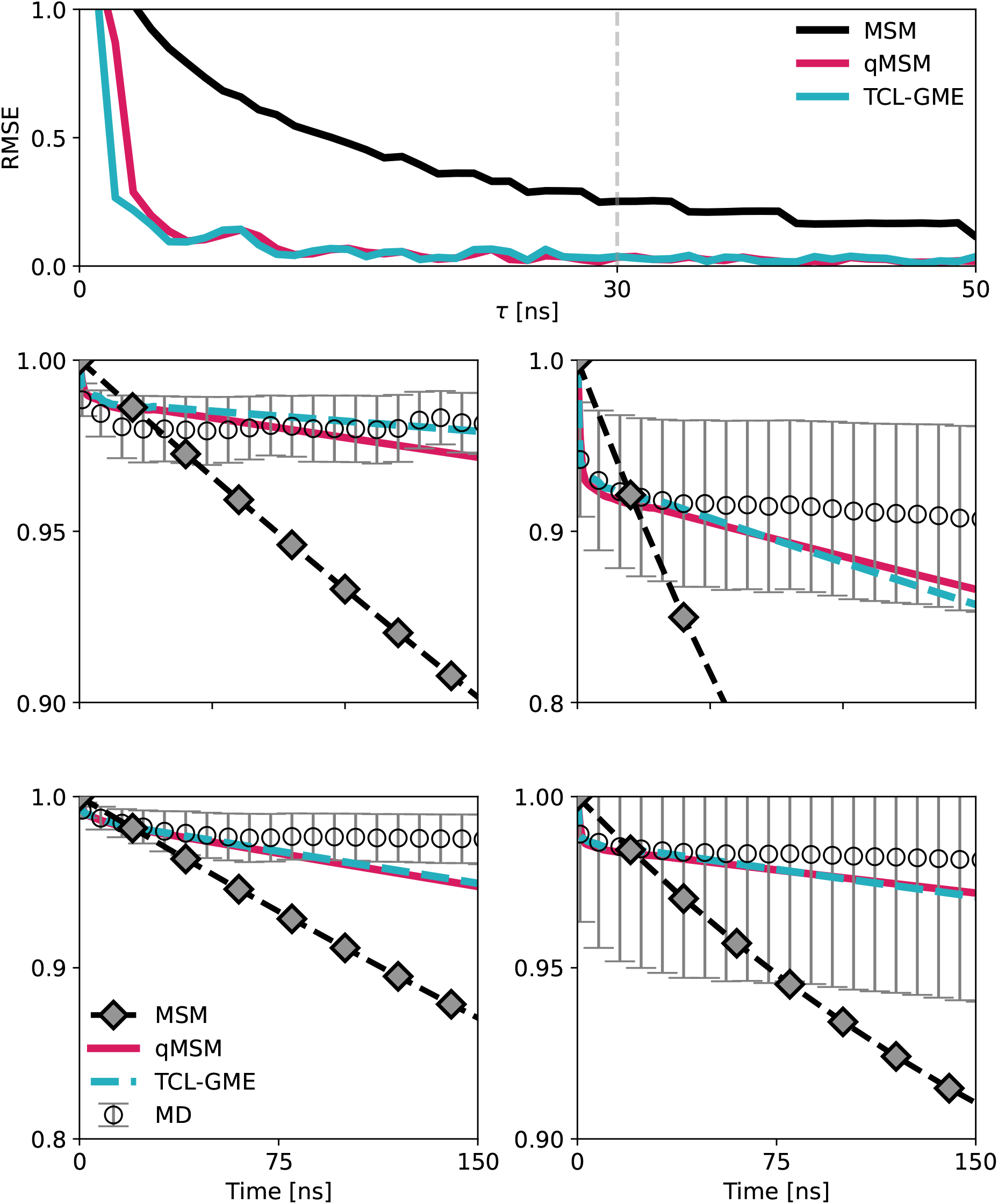
Illustrating the error associated with direct cutoffs of the memory kernel and time-local generator. (a) Root mean square error (RMSE) curve for the MSM, qMSM, and TCL-GME. The vertical line shows shows the error associated with a cutoff choice of (*τ*) of 30 ns. (b)-(e) Chapman-Kolmogorov test for the MSM, qMSM, and TCL-GME dynamics computed with *τ_L_* = *τ_K_* = *τ_R_* = 30 ns.

**FIG. 7.**
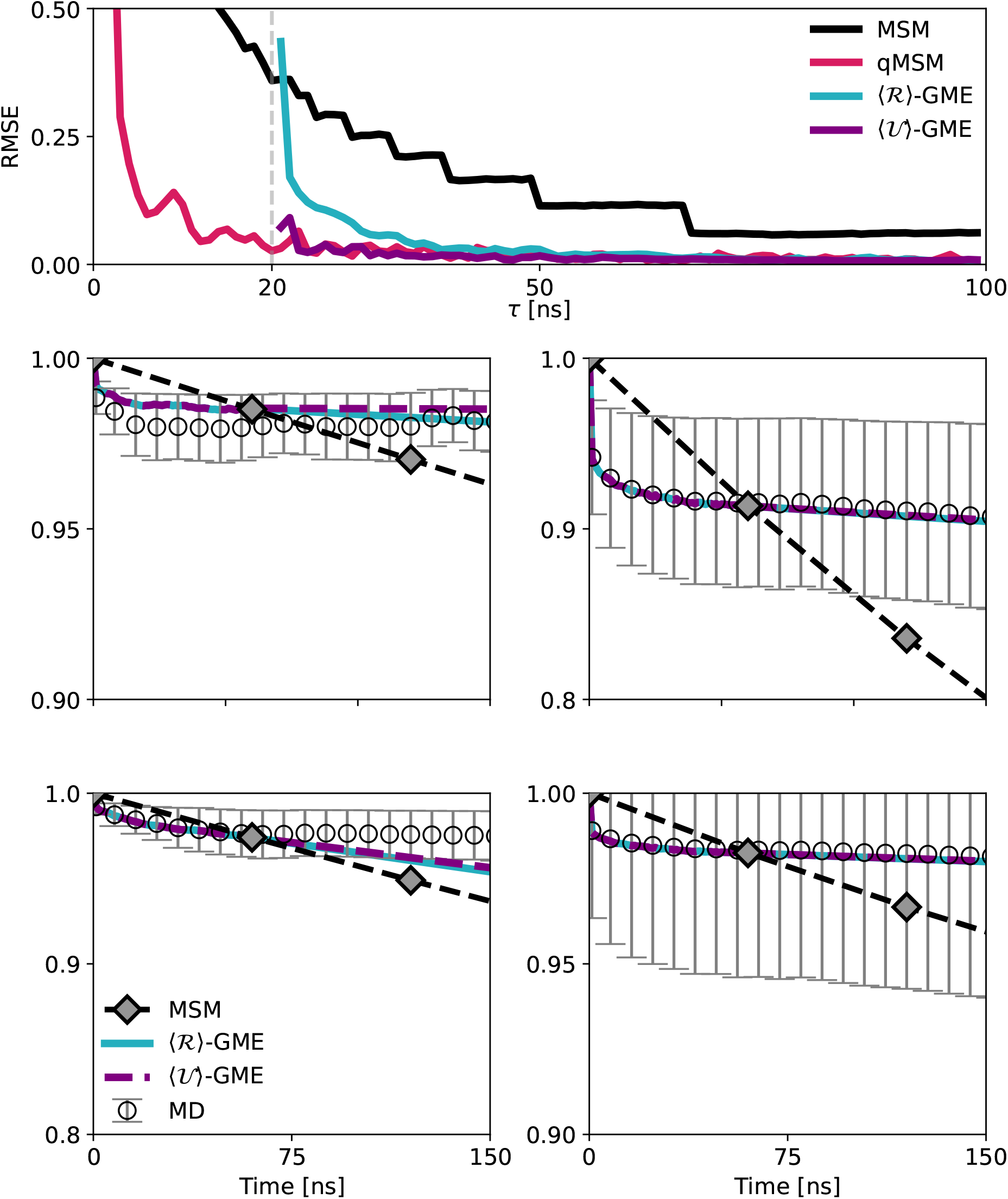
Convergence of the dynamics as predicted using the 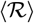-GME and 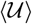-GME approaches. (a) Root mean square error (RMSE) curve for the MSM, 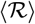, and 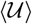-GME. The vertical line denotes the choice of *t_r_*, and thus the start of the averaging window. (b)-(e) Chapman-Kolmogorov test for the MSM, 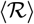, and 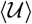-GME approaches with *t_r_* = 20 ns and *τ_R_* = 60 ns.

**FIG. 8.**
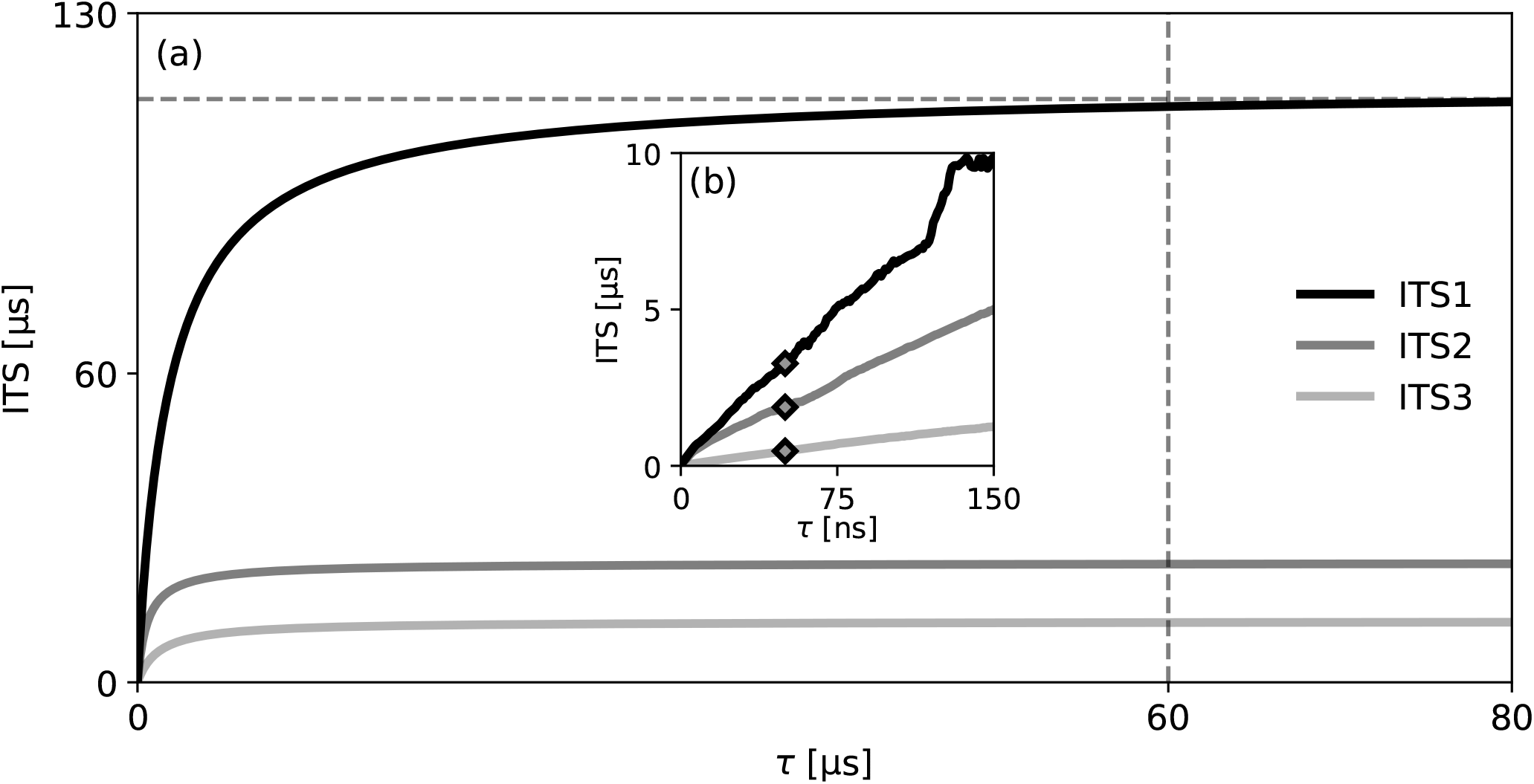
Extrapolating the MSM lagtime of the 4-state model of the human argonaute complex by employing the 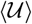-GME. (a) The 4-state implied time scales of computed with *τ_R_* = 50 ns, from which we determine the true MSM lag time to be ~ 60 μs. (b) The 4-state ITS computed from the reference dynamics, as displayed in Fig. 2 in the main text.

**FIG. 9.**
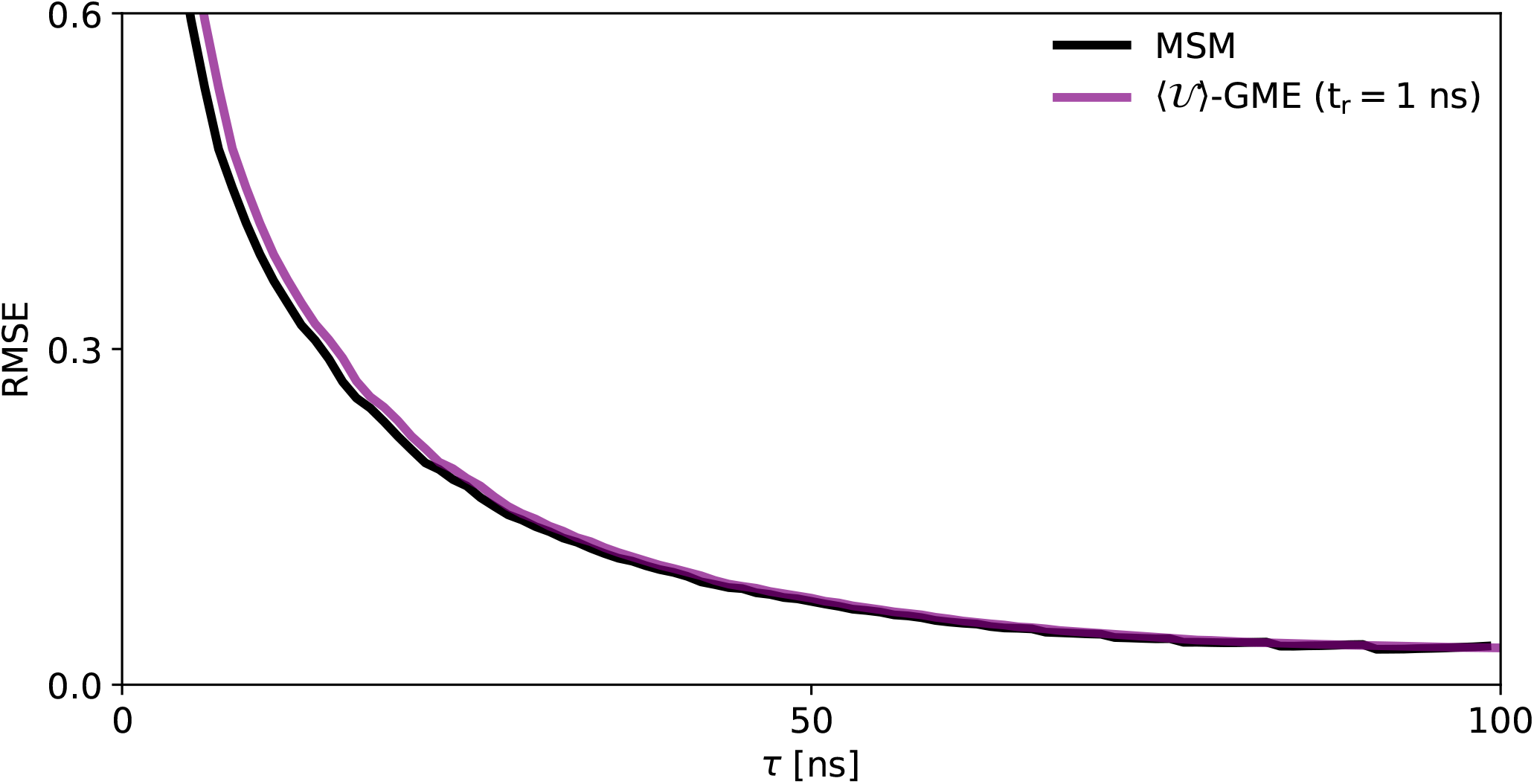
Illustrating that the errors, computed using an RMSE analysis, of calculating the 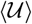-GME with *t_r_* = 1 ns and the MSM are roughly equivalent.

**FIG. 10.**
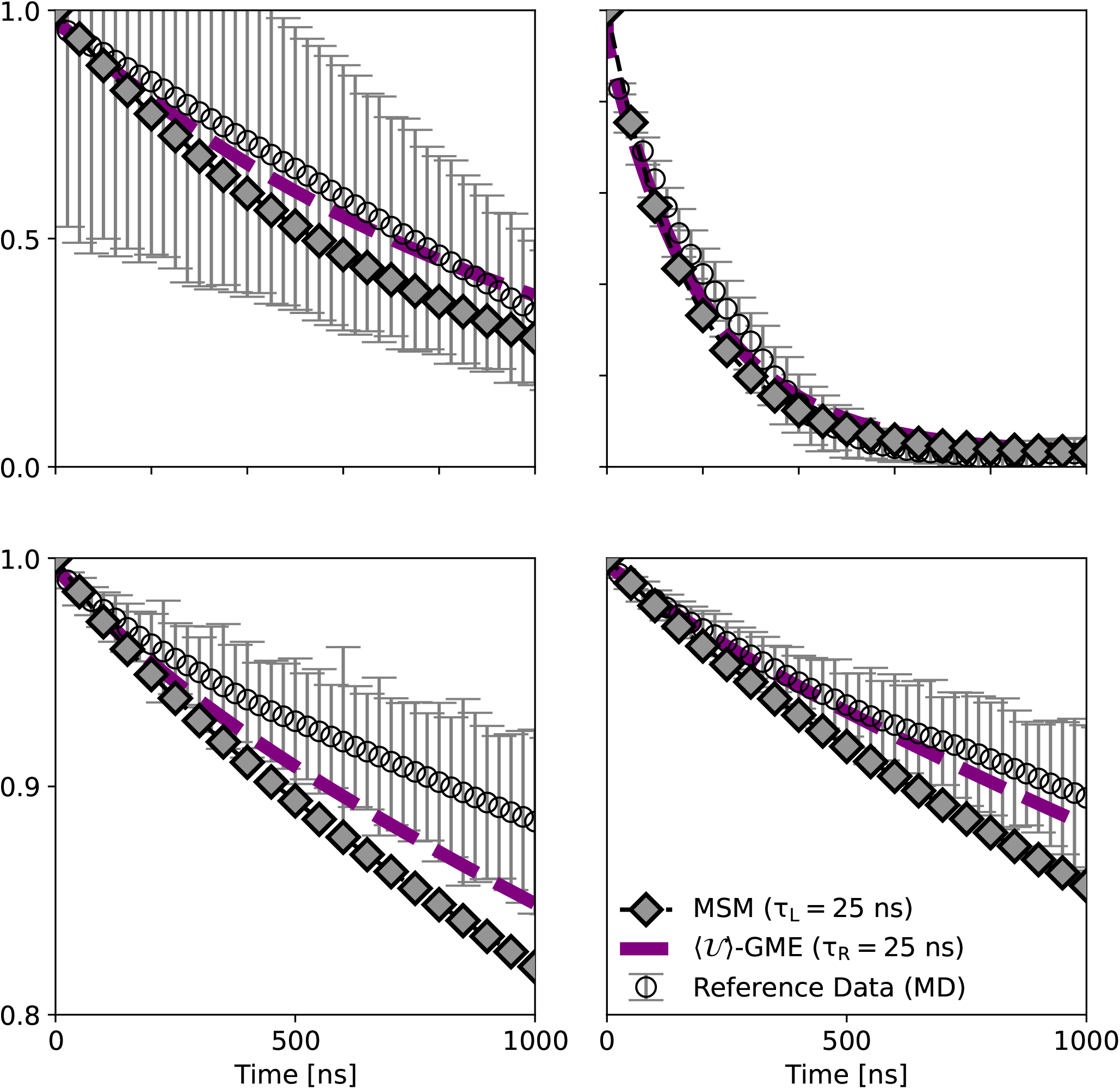
Chapman-Komolgorov test for the Chapman-Kolmogorov test for the MSM (*τ_L_* = 25 *ns*) and 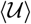-GME (*t_r_* = 20 ns; *τ_R_* = 5 ns)approaches.

**FIG. 11.**
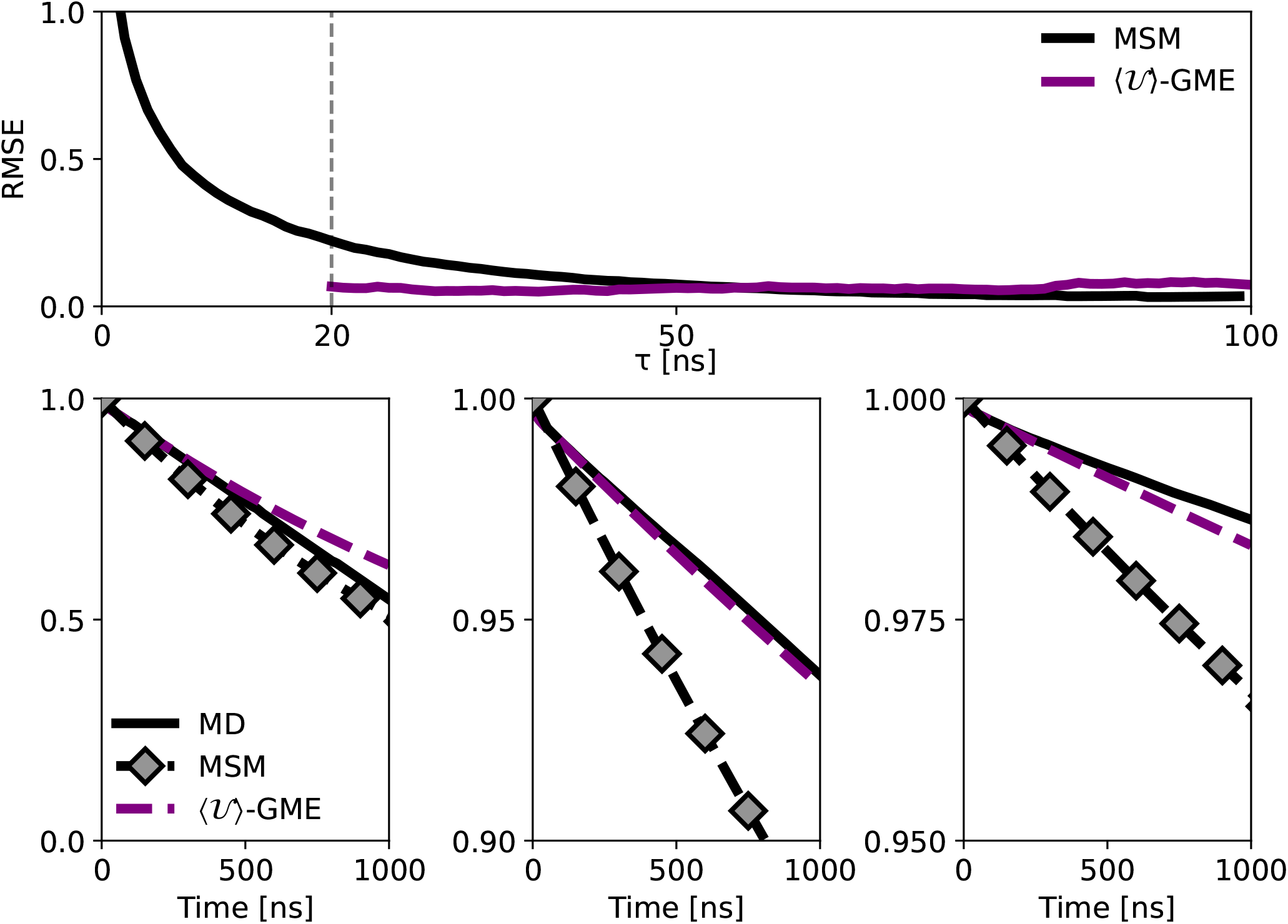
Illustrating the error associated with various cutoffs of the time-local generator to construct the 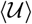 – *GME*. (a) Root mean square error (RMSE) curve for the MSM and 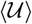-GME approaches (*t_r_* = 20 ns). The vertical line shows shows the error associated with a cutoff choice of (*τ*) of 50 ns. (b)-(e) Chapman-Kolmogorov test for the MSM, qMSM, and TCL-GME dynamics computed with *τ_L_* = *τ_K_* = *τ_R_* = 50 ns.

